# Plasticity of cold hardiness in the eastern spruce budworm, *Choristoneura fumiferana*

**DOI:** 10.1101/2021.04.02.438273

**Authors:** Skye Butterson, Amanda D. Roe, Katie E. Marshall

## Abstract

High latitude insect populations must cope with extreme conditions, particularly cold temperatures. Insects use a variety of cold hardiness mechanisms to withstand this temperature stress, and these can drive geographic distributions through overwintering mortality. The degree of cold hardiness can be altered by two evolved responses: phenotypic plasticity and local adaptation. Phenotypic plasticity can occur within or between generations (transgenerational plasticity; TGP), and local adaptation can evolve through directional selection in response to regional climatic differences. We used the eastern spruce budworm, *Choristoneura fumiferana* (Lepidoptera: Tortricidae) as a model to explore the role that variable winter temperatures play in inducing two aspects of plasticity in cold hardiness: TGP and local adaptation in phenotypic plasticity. This species is one of the most destructive boreal forest pests in North America, therefore accurately predicting overwintering survival is essential for effective management. While we found no evidence of TGP in cold hardiness, there was a long-term fitness cost to larvae that experienced repeated cold exposures. We also found evidence of local adaptation in both seasonal and short-term plasticity of cold hardiness. These findings provide evidence for the importance of phenotypic plasticity and local adaptation when modelling species distributions.

## Introduction

Cold hardiness during overwintering is an important determinant of temperate insect survival. Body fluids of small ectotherms are especially at risk of freezing in winter environments due to their relatively low thermal inertia (Lee, 2010). Freezing is deadly for most species; ice crystal formation causes mechanical damage to cells and cellular dehydration results in an accumulation of solutes and altered biochemical gradients (Lee, 2010). Among cold hardy insects, cold hardiness strategies can be divided into freeze tolerance and freeze avoidance (Sinclair, 1999). While freeze tolerant insects can survive internal ice formation, freeze avoidant insects survive low temperatures by depressing their supercooling point (SCP; the temperature at which their internal fluid freeze. The SCP is depressed through a number of processes such as: the dehydration of body water content (Han and Bauce, 1998), expression of low-molecular weight cryoprotectants like glycerol (Churchill and Storey, 1989), and production of ice binding proteins (IBPs; Duman, 2015).

The ability to respond to cold stress can vary by region, and these differences can arise through a combination of basal genetic adaptations and phenotypic plasticity (Sinclair et al., 2012). Adaptive differences arise as environmental conditions select for regionally adapted genotypes, leading to localized genetic differentiation. Phenotypic plasticity – the capacity of a single genotype to produce multiple environmentally-induced phenotypes – is an induced response to thermal variability (Pigliucci, 2001). These basal and plastic responses to cold stress may coevolve (Calosi et al., 2008), or manifest as a trade-off where populations with high basal cold hardiness have low phenotypic plasticity in variable thermal conditions (Kellermann et al., 2018). Disentangling these processes requires exploring cold hardiness among populations within a common-garden framework or using reciprocal transplant experiments (Overgaard et al., 2010).

Plastic responses to temperature can occur over multiple timescales. In the short term, an individual can develop resistance to stressful cold temperatures by hardening, which occurs rapidly after short exposures to sub-lethal conditions and is usually reversible (Kelty, 2007). Rapid cold hardening (RCH) is a form of hardening that occurs within minutes to hours after a cold exposure (Lee and Denlinger, 2010). Acclimation, on the other hand, is a more gradual adjustment to cold temperatures that occurs over a period of days to months (Lee, 2010). Both hardening and acclimation can lead to increased cold hardiness and may share biochemical mechanisms such as cryoprotectants and membrane modifications to achieve this phenotype, but there is evidence that they involve non-overlapping mechanisms (Rajamohan and Sinclair, 2009). Furthermore, repeated cold exposure can cause significant up-regulation of cryoprotectant concentrations beyond the effects of a single exposure (reviewed in Marshall and Sinclair, 2012). Responses to cold can depend on developmental stage and even age, with stage-specific tolerance shifting over time (Bowler & Terblanche 2008). For example, cold hardiness in *Drosophila melanogaster* varies across life stages (Jensen et al., 2008), with eggs having greater cold tolerance than other life stages. Plastic, stage-specific variation in temperature tolerance can contribute to mortality and population dynamics, which may be critical in determining the geographic distribution of a species (Bowler and Terblanche, 2008).

Plastic responses to cold stress can also extend beyond a single generation and impact future offspring through transgenerational plasticity. Maternal effects are the most frequently studied form of transgenerational plasticity (TGP; reviewed in Mousseau and Fox, 1998), but can also include paternal, grandparental, and potentially earlier generational effects. Transgenerational responses to temperature occur in many insect species (Woestmann and Saastamoinen, 2016). For example, diapause in the offspring of the cabbage beetle, *Colaphellus bowringi*, is modulated by the temperatures and photoperiods experienced by their parents. Moreover, the expression of diapause itself in the parental generation predicts the absence of diapause in the subsequent generation (He et al., 2018). Collectively, thermal conditions experienced by a population of insects can alter their immediate phenotypic response and have cascading impacts on the response and fitness of subsequent generations, making it critical to examine the phenotypic responses to cold stress over a range of timescales.

The ability to respond to temperature variability and the degree of that plastic response can, in itself, show local adaptation (Calosi et al., 2008; Kawecki and Ebert, 2004), resulting in variable expression among populations (Chown, 2001; Sinclair et al., 2012). For example, *Daphnia magna*, a widely distributed zooplankton, exhibits both local adaptation and adaptive plasticity in temperature tolerance (Yampolsky et al., 2013), with individuals from warmer sites expressing higher heat tolerance than those from cooler locations. The populations also showed variable degrees of acclimation to warmer temperatures that further increased heat tolerance, suggesting adaptive plasticity. This form of plasticity may be particularly important in species with large distributions, as climatic conditions can vary substantially along latitudinal or altitudinal gradients, providing a mechanism for local populations to respond to regional temperature conditions.

The eastern spruce budworm (*Choristoneura fumiferana* Clemens, 1865) is a destructive forest pest found throughout the northern boreal forest (Fig. 1), whose distribution spans ~24 degrees of latitude and over 80 degrees of longitude. This distribution represents a large winter temperature gradient, where populations in the southern range will experience a much milder winter climate than those in the northern latitudes. Spruce budworm survives winter as second instar larvae in silken overwintering structures (called hibernacula) and undergoes a six to eight month dormancy (Han and Bauce, 1993; Marshall and Roe, n.d.; Régnière, 1990). Ice formation is lethal to *C. fumiferana*, so the SCP represents a physiological and fitness limit for this freeze avoidant species. Second instar larvae are the primary overwintering stage and can depress their SCP to ~-35°C (Han and Bauce, 1995a) by accumulating ~ 0.8 M glycerol (Han and Bauce, 1995b), expressing IBPs (Qin et al., 2007; Tyshenko et al., 1997), and decreasing their internal water content (Bauce E and Han, 2001; Han and Bauce, 1998; Marshall and Roe, n.d.). While capable of surviving very brief exposures to temperatures near their SCP, exposure to −15 °C for more than ten days can result in mortality (Han and Bauce, 1995a). There is also evidence that fluctuating temperatures can further induce significant plasticity in cold tolerance on short time scales (*i.e*. days, Marshall and Sinclair, 2015), even in seasonally-acclimated individuals. *C. fumiferana* larvae overwintered outside have significantly lower SCPs than larvae overwintering in laboratory conditions (Han and Bauce, 1993), suggesting that fluctuating conditions may be a critical cue for inducing a greater response to cold stress than expressed under constant temperatures.

**Fig. 1.**
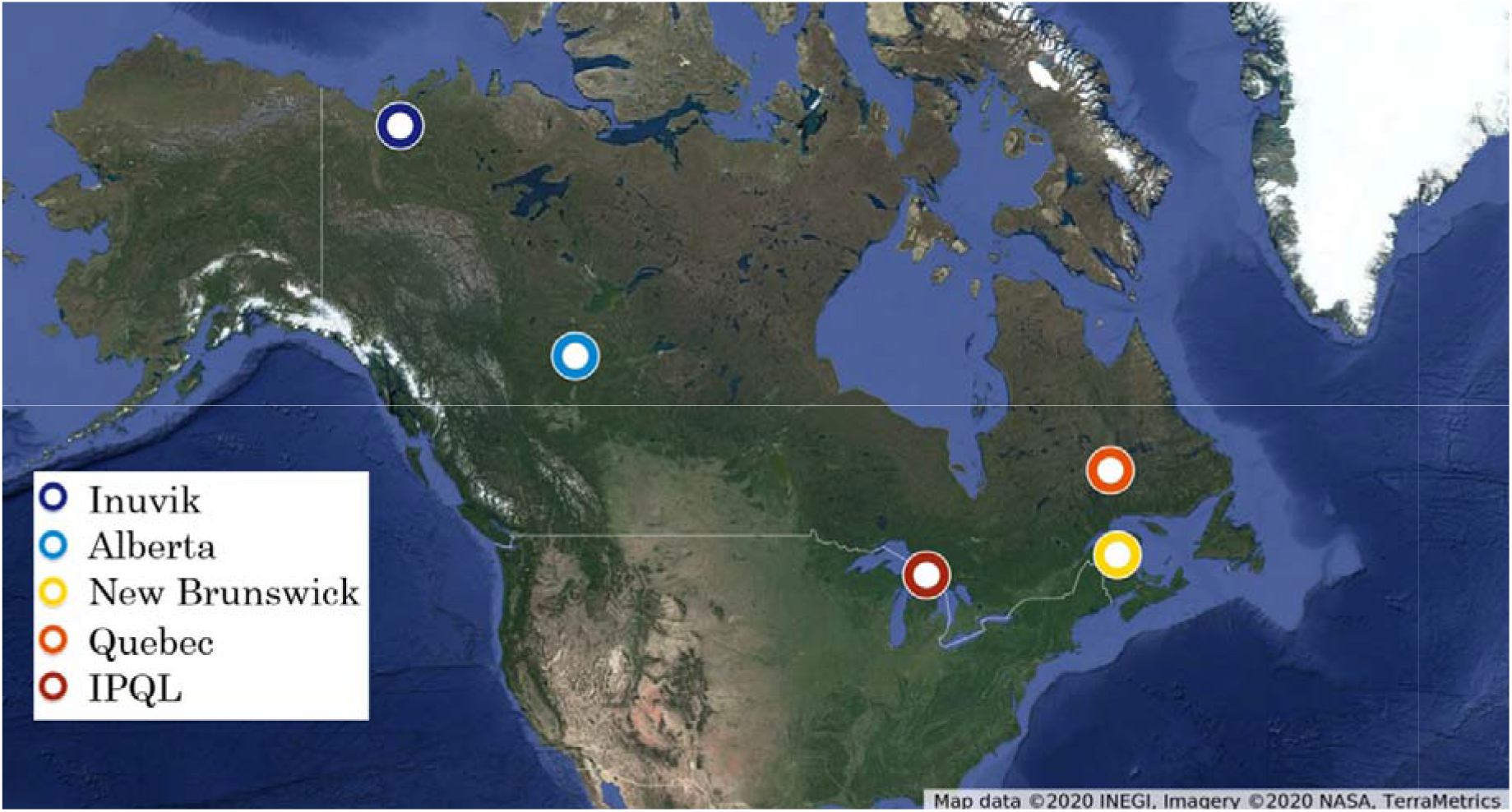
Map of population sampling localities.

The vast majority of cold tolerance work for *C. fumiferana* has been based on a laboratory strain maintained at the Insect Production and Quarantine Laboratories (IPQL, Natural Resources Canada, Great Lakes Forestry Centre, Sault Ste. Marie, Ontario, Canada). The IPQL laboratory strain was established from multiple populations in Ontario and has been in culture for over 60 years (Roe et al., 2017). Genetic drift and historical population bottlenecks, as well as potential laboratory adaptation to constant temperatures, could limit the general applicability this strain to characterize cold stress responses of wild populations. The ability of *C. fumiferana* to withstand low temperature stress, particularly in northern latitudes, is crucial to understanding their overwintering capacity, and by extension, their geographic distribution (Gray, 2008; Régnière, 2009). Currently, winter conditions at the northern edges of their range exceed these published limits, which suggests that cold hardiness capacities in the eastern spruce budworm may be significantly under-reported and vary throughout their range. Hence, we need to explore the intraspecific variation in basal and plastic cold tolerance to determine the overwintering capacity of the spruce budworm.

Spruce budworm larvae do not feed after hatching, but prepare for diapause immediately following eclosion. This means that the energy reserves needed to survive winter and initiate spring development are entirely dependent on parental investment (Sanders, 1991). Therefore, conditions or stress experienced by the parental generation could affect the cold tolerance of their offspring via TGP. For example, acclimating *Calliphora vicina* to lower fall temperatures decreases the SCP of their offspring by 3 °C (Coleman et al., 2014). In *C. fumiferana*, previous work has found transgenerational fitness effects on offspring fitness, driven by parents’ nutritional status as sixth-instars (Carisey and Bauce, 2002). It is, however, unclear whether TGP influences the overwintering capacity of *C. fumiferana* or contributes to its adaptive plasticity in cold tolerance.

The basic mechanisms of cold hardiness in *C. fumiferana* are well understood, however, we know little about the role plasticity and local adaptation plays in the survival of this widespread species. Using new laboratory strains of *C. fumiferana* (Perrault et al., 2021), we tested two hypotheses: 1) that spruce budworm populations have local adaptation in the plasticity of cold hardiness on either seasonal or short-term timescales, and 2) that this plasticity induces TGP in cold hardiness. We found that while fluctuating temperatures induced plasticity in cold hardiness, there was no evidence that this plasticity was transgenerational. However, strains of *C. fumiferana* from higher latitudes exhibited greater basal cold hardiness and significantly more plasticity than those from lower latitudes.

## Materials and methods

### Experimental animals

We performed all experiments on second-instar diapausing *Choristoneura fumiferana* caterpillars obtained from IPQL. Transgenerational plasticity experiments were conducted on the “IPQL” strain (Roe et al., 2017) established in 1961 (Glfc:IPQL:Cfum). Local adaptation in plasticity experiments were conducted on colonies established from wild populations as reported in Perrault et al. (2021; Fig. 1). Regional colonies used in this study originated from Campbellton, New Brunswick (Glfc:IPQL:CfumBNB01, “New Brunswick” 47°59’13.1”N 66°40’37.8”W), Manic-Cinq, Quebec (Glfc:IPQL:CfumMQC01, “Quebec” 52°51’07.1”N 67°06’35.5”W), High Level, Alberta (Glfc:IPQL:CfumHAB, “Alberta” 58°50’71” N 117°14’03” W), Inuvik, Northwest Territories (Glfc:IPQL:CfumINT01, “Inuvik” 68°21’56.2”N 133°42’04.9”W). The Alberta and Inuvik populations correspond to the “Central” genomic cluster identified in Lumley et al. (2020), with the remaining populations contained within the “Eastern” cluster. These regional strains were kept in culture for one (late diapause measures) or two generations before use (all other measures). We received dormant larvae in newly-spun hibernacula embedded in gauze from IPQL via overnight shipping on ice packs to the University of British Columbia. Once received, larvae were placed into an incubator (MIR-154, Sanyo, Bensenville, USA) held at 2 °C in constant darkness.

### Experimental design: local adaptation in phenotypic plasticity

We assessed local adaptive plasticity in four colonies of eastern spruce budworm recently derived from wild populations (Inuvik, Alberta, Quebec, New Brunswick). We tested two aspects of locally adaptive plasticity in cold hardiness during diapause: 1) in constant dormancy temperatures, and 2) following variable cold exposure regimes. The first was tested by measuring larvae either six or twelve weeks into diapause (held at 2 °C under constant darkness), constituting the “early” and “late” diapause groups, respectively. To test the effect of variable temperatures, early diapause groups were further separated into “basal” or “inducible”. Individuals in the “inducible” treatment group were held at 2 °C for six weeks, then exposed to five repeated cold events of −15 °C for 12 hours with a return to 2 °C for 12 hours (ramp rate of 0.05 °C/minute). No caterpillars from the New Brunswick population were exposed to the inducible treatment due to limited sample numbers.

### Experimental design: Transgenerational plasticity

We explored whether parental cold stress would generate a transgenerational response in *C. fumiferana* larval cold tolerance. First, we exposed the parental population to different temperature regimes, then measured SCP, biochemical properties, and lower lethal temperature (LLT). We exposed 980 diapausing IPQL caterpillars to four cold temperature regimes: control, single 1, single 2, and repeated. “Control” insects were kept at 2°C for six weeks. “Single 1” insects were kept at 2 °C for six weeks, then exposed to −10 °C for 12 hours (including a ramp rate of 0.05 °C/minute) and measurements were made after 24 hours at 2 °C. “Single 2” insects were kept at 2 °C for six weeks, then exposed to −10 °C for 12 hours (including a ramp rate of 0.05 °C/minute) and measurements were made after 152 hours (~ 6 days and 8 hours) at 2 °C to account for differences in the time since first cold exposure between “single 1” and “repeated”. “Repeated” insects were kept at 2 °C for six weeks, then exposed to five repeated cold events of −10 °C for 12 hours with a return to 2 °C for 12 hours (ramp rate of 0.05 °C/minute). We provide detailed descriptions of these cold exposure techniques below (see *Cold exposures*). All temperature treatments were conducted in total darkness. Following temperature exposures, we haphazardly selected 20 individual larvae for SCP measurements, five sets of 10 larvae for biochemical assays, and five sets of 20 larvae for LLT measurements. We also retained a subset of individuals (10 sets of 10 larvae) from the parental treatment groups to generate F1 progeny and repeated the cold tolerance measurements on the F_1_ progeny (i.e. SCP and biochemical properties).

### Cold exposures

To conduct cold exposures, we placed microcentrifuge tubes containing larvae into a milled aluminum block (Sinclair et al., 2015) connected to a programmable refrigerated circulating bath (Lauda Proline RP 3530, Wurzburg, Germany) containing 50:50 ethylene glycol:water. Ten 36 AWG Type T (copper-constantan) thermocouples (Omega Engineering Inc., Laval, Canada) were placed in the block to monitor temperature. These thermocouples were interfaced with PicoTech TC-08 thermocouple interfaces connected to a computer running PicoLog software (Pico Technology, Cambridge, U.K.) taking temperature samples in the block every 0.5 seconds. The bath was set to cycle between 2 and −15 °C (OR −10 °C for TGP exposures) for 12 hours each including ramping rates between these temperatures of 0.05 °C/min for five full cycles. Larvae were returned to the incubator in diapausing conditions for 24 hours to recover after the exposure.

### Measures of cold hardiness

#### Supercooling point (SCP)

SCPs were measured as in (Strachan et al., 2011). Briefly, we removed larvae from their hibernacula and stored them in 1.8 mL microcentrifuge tubes until SCP measurements. We attached 20 individual caterpillars to 36 AWG Type T copper-constantan thermocouples with a thin layer of vacuum grease. We threaded these thermocouples through the pierced top of the 1.8 mL microcentrifuge tubes and held in place with adhesive putty.

We floated each tube on a 60:40 methanol: water solution in a programmable refrigerated circulating bath (Lauda ECO RE 1050, Wurzburg, Germany). The refrigerated circulating bath was cooled from 2 to −38 °C at a rate of 0.09 °C/minute. SCP was recorded as the temperature immediately prior to the onset of the exotherm (Lee, 2010; Sinclair et al., 2015). Late diapausing Inuvik caterpillars could not be reliably frozen using the abovementioned method. In this case, SCPs were estimated by placing computer-interfaced thermocouples attached to individual caterpillars in microcentrifuge tubes in Styrofoam freezer boxes and then into a −80 °C freezer. The cooling rate for this exposure can only be estimated in this case as 9 °C/minute.

### Biochemical assays for energetics and metabolites

We removed larvae from their hibernacula and stored them as in SCP treatment. We then homogenized groups of 10 caterpillars with approximately 90 0.5 mm Zirconium oxide beads (Next Advance Inc., Averill Park, USA) in a Bullet Blender (Storm 24, Next Advance Inc., Averill Park, USA) for two minutes at the highest speed. After initial blending, we added 50 μL 0.05% Tween 20 and repeated blending. An additional 250 μL 0.05% Tween 20 was added and the sample was mixed using a vortexer (Vortex-Genie 2, Scientific Industries Inc., Bohemia, USA). Following mixing, we centrifuged each sample (Allegra 64R, Beckman Coulter Canada Inc., Mississauga, Canada) for 10 min at 15,000 × *g*. We removed two aliquots of the supernatant and stored the samples at −80 °C for later assays.

Glycerol, glucose, glycogen and protein content was measured using spectrophotometric assays following (Marshall and Sinclair, 2015) using glycerol, glucose and Type II glycogen from oyster and bovine serum albumin as standards, respectively. Briefly, glycerol was measured using a Free Glycerol kit (MAK117, Sigma-Aldrich Canada Co., Oakville, Canada). Glucose was measured using a hexokinase-based Glucose assay kit (GAHK20, Sigma-Aldrich Canada Co.). Glycogen content was measured using the same kit following an 8 h amyloglucosidase (A9228, Sigma-Aldrich Canada Co.) digestion in a dark drawer at room temperature. Soluble protein was measured using a Bicinchronicinc acid kit (BCA1, Sigma-Aldrich Canada Co.). Absorbance of each reaction was measured in a spectrophotometer (Spectra Max M2, Molecular Devices, San Jose, USA) and calculated concentrations are reported in μmol/individual.

### Additional measures for TGP experiment: Lower lethal temperature (LLT) and rearing F_1_ progeny

To estimate LLT, five groups of 20 caterpillars within their hibernacula from each treatment group were exposed to either −15, −20, −25, −30, or −35 °C, as described in (Sinclair et al., 2015). Groups of larvae were exposed to their corresponding temperature treatment for 4 hours, by putting the pierced 1.8 mL microcentrifuge tubes into a milled aluminum block as described for the cold exposure treatments. After exposure, they were transferred to Petri dishes. Mortality was assessed under the microscope (MEB126, Leica, Wetzlar, Germany) by checking if caterpillars were dehydrated, out of their hibernaculum or immobile, at five different time points throughout development (one week after exposure, at the end of diapause (20 weeks from the onset of diapause), at thinning (10 days after ending diapause, between instar 3 and 4) and at pupation, following the rearing protocols explained below. Dead caterpillars were removed at each time point. The temperature at which 50% of the group survived (LT_50_) was calculated at each time point by assessing mortality to determine both change in LT_50_ after different cold exposures (treatments) and change in LT_50_ over development (time).

After the cold exposures, the mating group was returned to diapausing conditions (2 °C in constant darkness) for a total of 20 diapausing weeks (including the initial six weeks pretreatment and treatment time). Following this, they were put into feeding/developing conditions (24 °C, with a 16:8 light: dark cycle) as detailed in (Marshall and Sinclair, 2015). Briefly, groups of 10 caterpillars were put onto diet cups with artificial diet (McMorran, 1965) purchased through Insect Production and Quarantine Services (https://www.nrcan.gc.ca/science-data/research-centres-labs/forestry-research-centres/great-lakes-forestry-centre/insect-production-services/13467). Insects were placed on fresh diet once a week to avoid the accumulation of frass, microbes and mold. Ten days later, one caterpillar was put into each cup (thinning). At pupation, we sterilized pupae with a 10% bleach solution under a fume hood, rinsed with deionized water and left to dry on paper towels. They were then sexed under a microscope and the first 40 individuals were weighed on a microbalance (CP 124 S, Sartorius, Göttingen, Germany). Males and females were moved to separate ventilated plastic containers to emerge. These emergence chambers were checked daily for emerged adult moths. The first 20 male and female moths (total of 40 moths) that emerged were put into 20 L clear plastic bags with three sets of five 5 × 5 cm strips of waxed paper stapled together. A 50:50 female and male ratio was maintained in each bag. Each treatment’s mating chamber was sprayed with deionized water before and after take-down. The three mating chambers were kept in environmental chambers (23 ± 3°C, 55 ± 10% RH, 16L:8D) in the Faculty of Forestry at the University of British Columbia. Mating chambers were set-up for a week, after which moths died and egg masses were collected.

Emergence pans for egg masses were set-up using 30 × 25 × 5 cm baking trays lined with gauze and sealed with Parafilm and electrical tape to avoid larval escape. Once second-instar caterpillars began spinning their hibernacula, the emergence pans were opened to count the number of larvae that had emerged and spun their hibernacula. The gauze was then removed, wrapped it with Parafilm, and put it in an incubator held at 2 °C in constant darkness.

After six weeks into diapause, all measures of cold hardiness were repeated with F1 second-instar larvae.

### Statistics

All statistical tests and data plots were conducted in RStudio (version 1.1.463, 2018). To test for transgenerational plasticity, a Type II ANOVA model was fitted for each cold hardiness measure using the aov function in the car package (Fox and Weisberg, 2019) with generation and treatment as predictors. To test for local adaptation in cold hardiness, the same model was fitted with population, diapause stage and treatment as predictors. For biochemical assays, we used protein concentration as a covariate in the model. LLTs were calculated at each assessed time point using a generalized linear model with binomial error distribution and the dose.p function with p set to 0.5 from the MASS package (Venables and Ripley, 2003). Alpha was set to 0.05 for all tests. p-values less than 0.01 are reported as such. All significant interactions were further investigated using TukeyHSD posthoc tests.

## Results

### Local adaptation in phenotypic plasticity of cold hardiness

Spruce budworm populations showed significant differences in cold hardiness, particularly as larvae progressed in diapause. Late diapause individuals from the Inuvik population had a significantly lower SCP compared to other populations and times tested (population: F_(2,119)_=4.87, p<0.001; time: F_(1,119)_=16.06, p<0.001; population × time: F_(2,119)_=5.43, p<0.01; Fig. 2A). Population had a significant effect on basal glycerol content (population: F_(2,29)_=13.85, p<0.001; Fig. 2B). In this case, larvae from Inuvik significantly increased their glycerol by the late diapause time point, whereas larvae from New Brunswick significantly decreased glycerol content during the same timeframe. Glycogen concentrations were significantly affected by time (time: F_(1,29)_=11.27, p<0.01; Fig. 2C). Population and time in diapause also had a significant effect on basal total carbohydrate concentrations (population: F_(2, 29)_=3.196, p=0.020; time: F_(1, 29)_=15.96, p<0.001), with New Brunswick larvae in early diapause having lower total carbohydrate content than the early diapause larvae from Inuvik and Quebec (Fig. A2).

**Fig. 2.**
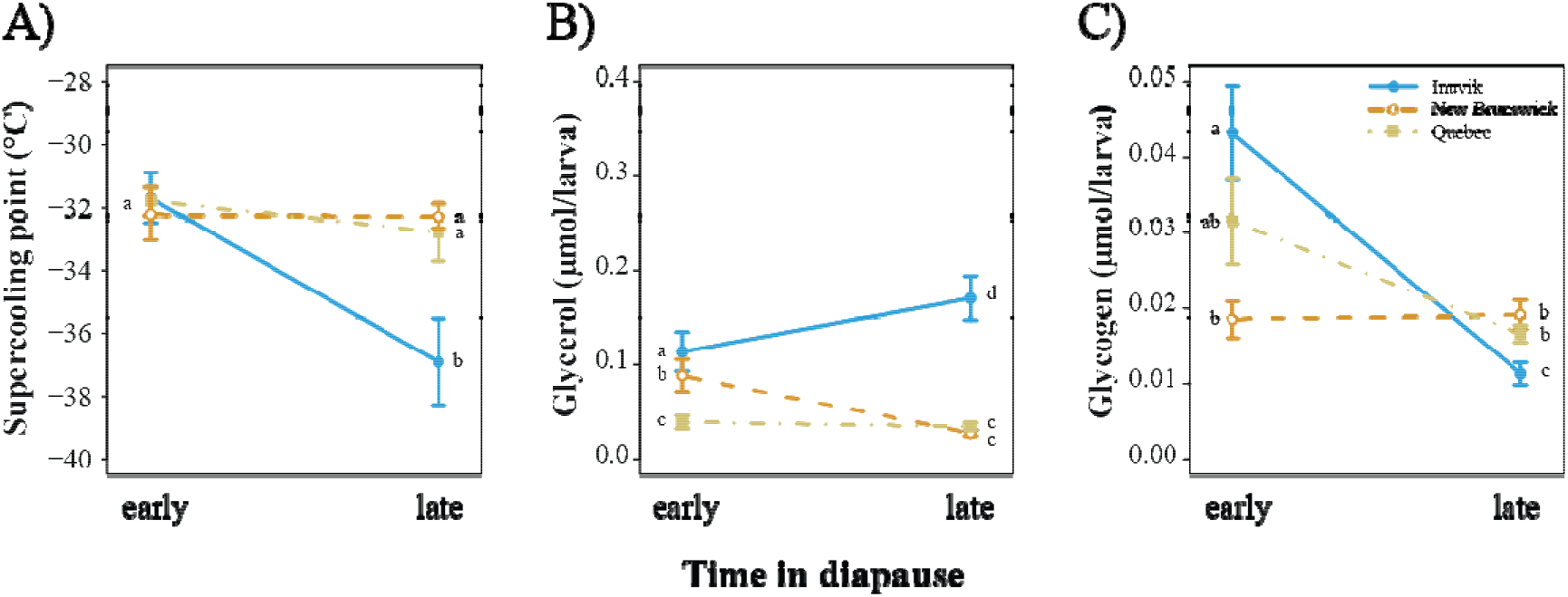
Testing for local adaptation in seasonal plasticity without pretreatment. A) Supercooling point (SCP; °C), B) glycerol per larva (μmol) and C) glycogen per larva (μmol) of second-instar *Choristoneura fumiferana* across different populations measured at either early (6 weeks) or late (12 weeks) into diapause. Glycogen is measured in glucose units. Points with the same letter are not statistically significantly different in an ANCOVA with protein mass as a covariate.

We exposed larvae to repeated cold events to examine the differences between basal cold hardiness and an inducible plastic response. We found a significant interaction between population and repeated cold exposures on SCP (population: F_(3,169)_=10.32, p<0.001; treatment: F_(1,169)_=15.80, p<0.001; population × treatment: F_(3,169)_=3.654, p=0.014; Fig. 3A). A significant effect of population and treatment was found on glycerol concentrations (population: F_(3,39)_=36.47, p<0.001; treatment: F_(1,39)_=31.31, p<0.001; population × treatment: F_(3,39)_=7.24, p<0.001; Fig. 3B). Alberta and Inuvik decreased their SCP after exposure, whereas Quebec and the IPQL remained the same. For glycogen; there was no effect of population or treatment (population: F_(3,39)_=0.123, p=0.12; treatment: F_(1,39)_=0.534, p=0.47; Fig. 3C). The same was true for total carbohydrate, there was no significant effect of population or treatment (population: F_(3,39)_=2.765, p=0.058; treatment: F_(1,39)_=0.88, p=0.35; Fig. A3).

**Fig. 3.**
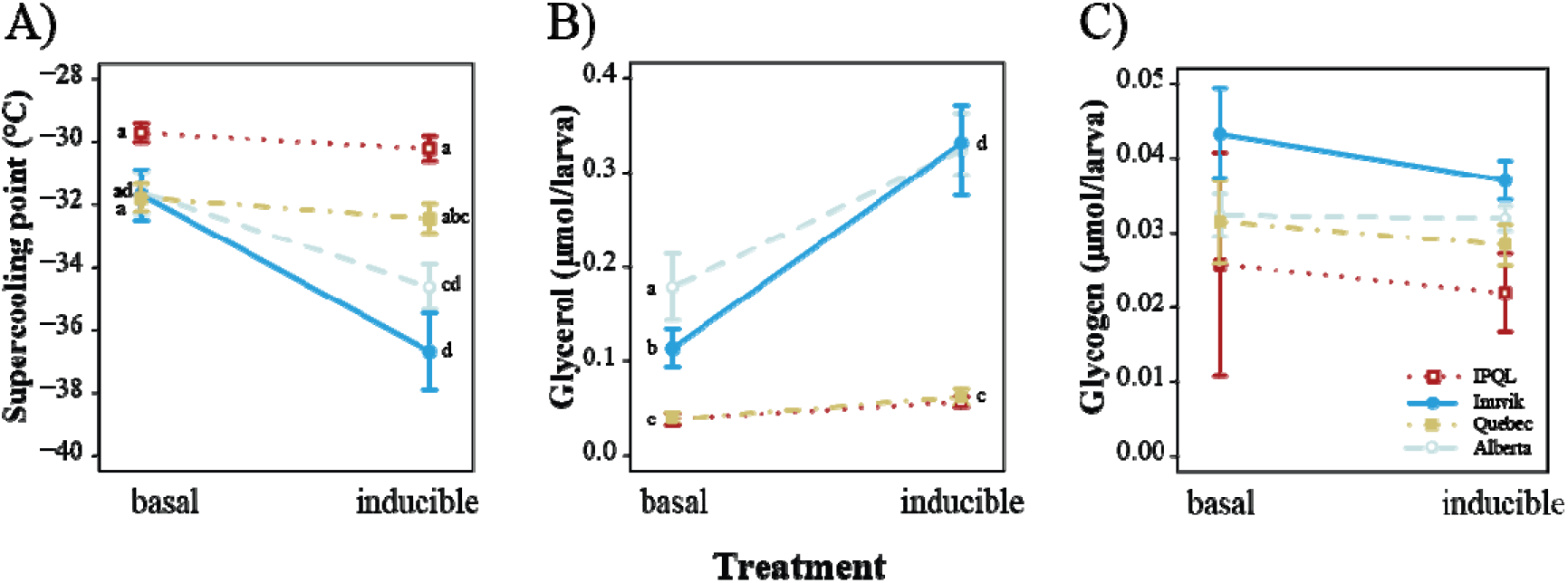
Testing for local adaptation in short-term response to temperature variation. A) Supercooling point (SCP; °C), B) glycerol per larva (μmol) and C) glycogen per larva (μmol) of second-instar *Choristoneura fumiferana* before (“basal”) and after (“induced”) five exposures to −15 °C. Different letters indicate statistically significant comparisons (p≤α). All exposures were conducted on early diapause larvae. Glycogen is measured in glucose units. Points with the same letter are not statistically significantly different in an ANCOVA with protein mass as a covariate.

### Transgenerational plasticity of cold hardiness

In the IPQL strain, repeated cold exposures did not significantly change SCP of either generation, although SCP was higher in the F_1_ generation (generation: F_(1,139)_=63.77, p<0.01; Fig. 4).

**Fig. 4.**
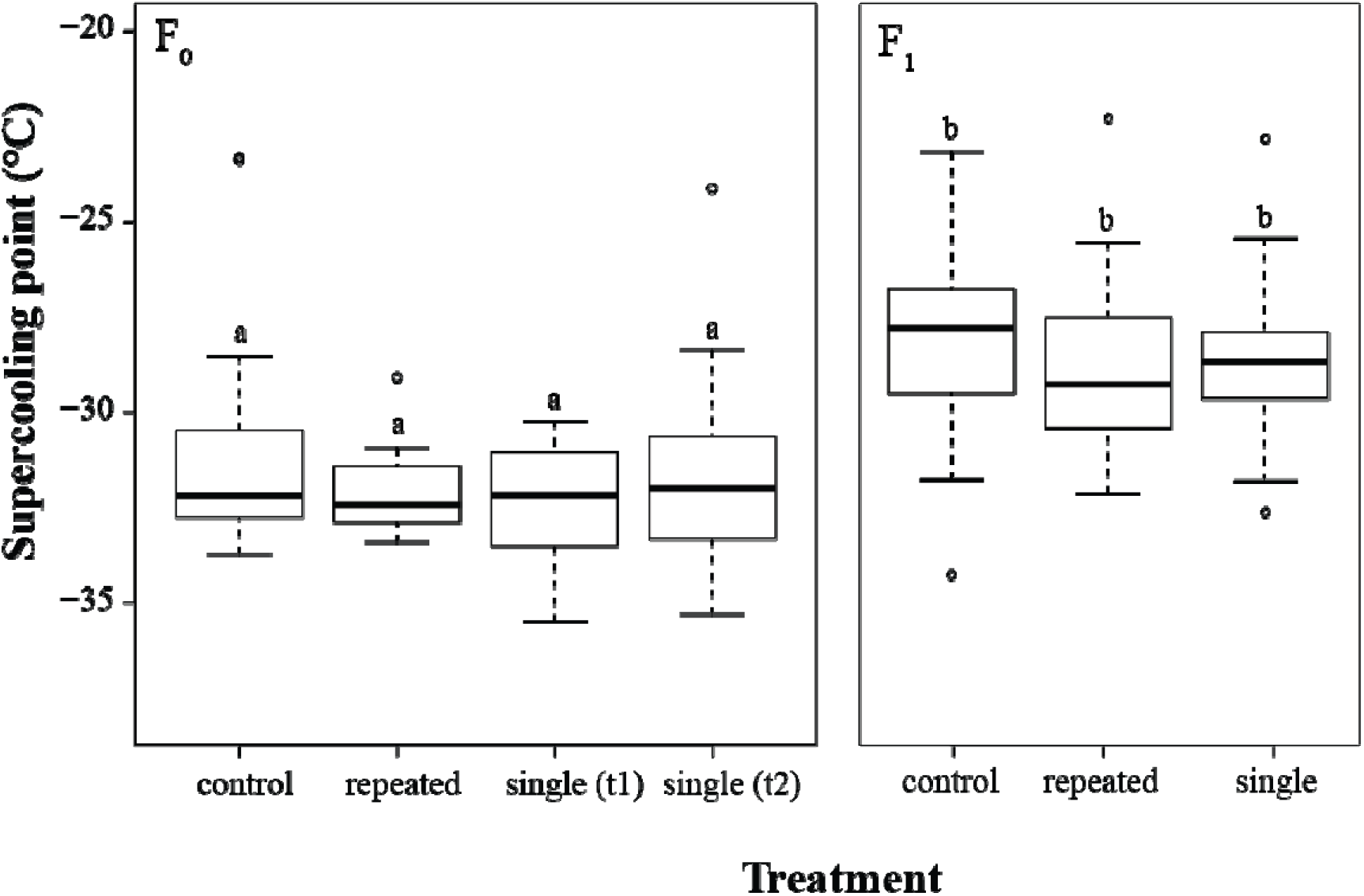
Supercooling points (SCP) (°C) as a result of cold exposures and generation in second instar IPQL *Choristoneura fumiferana* larvae. Bold line inside box shows median, lower and upper box boundaries show 25th and 75th percentile, respectively, lower and upper error lines show 10th and 90th percentile, respectively. Treatment groups are Single T1 (larvae exposed to a single cold exposure of −10 °C for 12 hrs and assessed 24 hours after), Single T2 (larvae exposed to a single cold exposure of −10 °C for 12 hrs and assessed at the same time as the repeated group) and Repeated (larvae exposed to five cold exposures of −10 °C for 12 hrs and assessed 24 hours after). Different letters indicate statistically significantly different comparisons (p≤α).

We calculated LT_50_ for each treatment group at four time points following exposure to five temperatures. We assessed mortality one week after exposure, at the end of diapause (20 weeks from the onset of diapause), thinning (between instar 3 and 4) and following pupation (Fig.s A4-7). The effect of cold exposure on LT_50_ depended on the time of assessment and treatment (time point: F_(3,15)_=58.63, p<0.001; treatment: F_(3,15)_=4.18, p=0.047). Individuals exposed to a single cold event (but sampled at the same time as the repeated group, Single 2) had a higher LT_50_ at every time-point assessed, except at the end of diapause, compared to other treatments (Fig.. 5). Additionally, LT_50_ increased with time as mortality effects accrued through development. Larvae that experienced repeated cold exposures had the lowest LT_50_ at every time point.

**Fig. 5.**
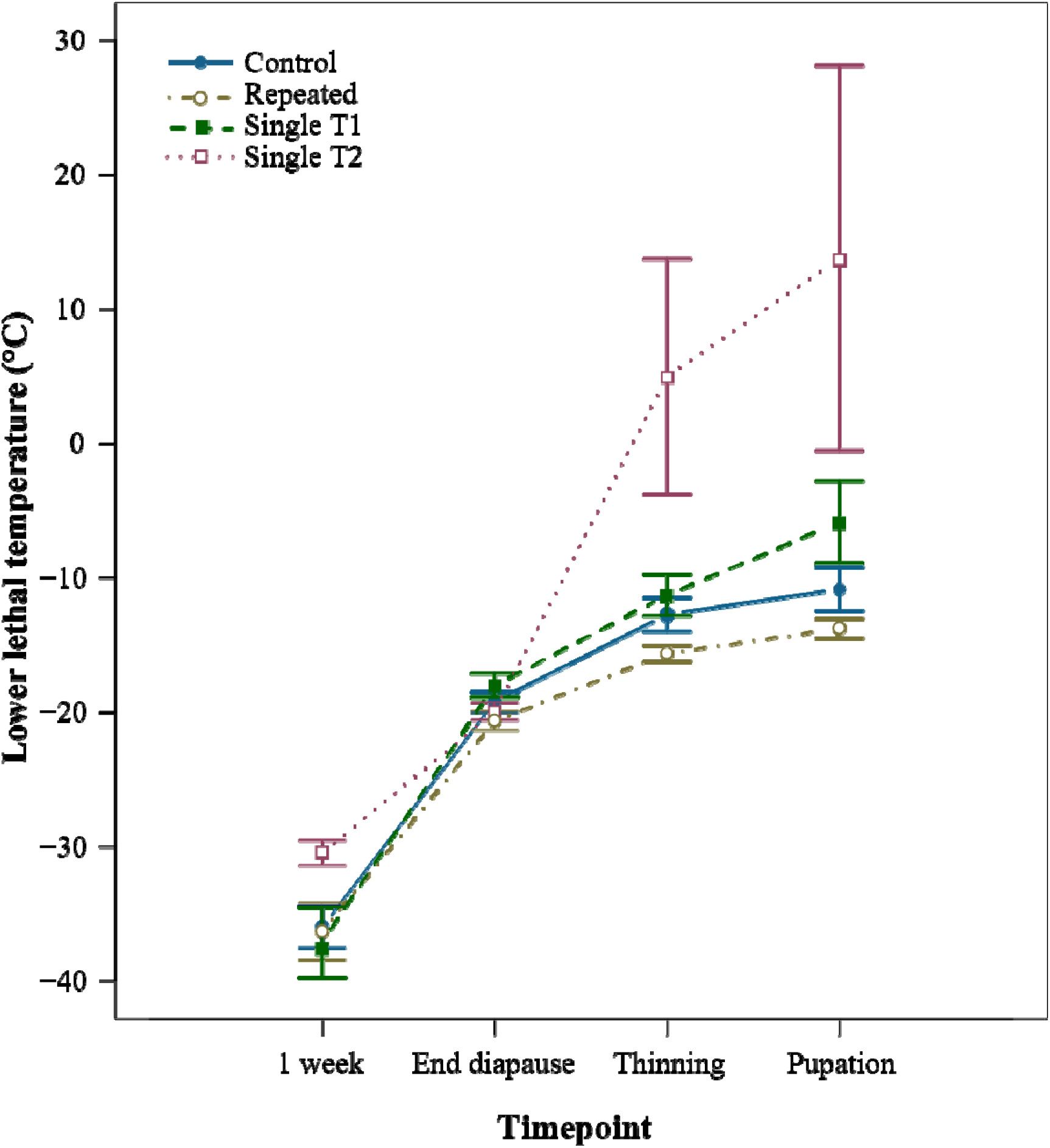
Lower lethal temperature (LLT), displayed as LT_50_, of F_0_ second-instar IPQL (*Choristoneura fumiferana*) larvae after different cold exposures shown in color. Error bars show standard error of the LT_50_ estimate. Treatment groups are Single 1 (larvae exposed to a single cold exposure of −10 °C for 12 hrs and assessed 24 hours after), Single 2 (larvae exposed to a single cold exposure of −10 °C for 12 hrs and assessed at the same time as the repeated group) and Repeated (larvae exposed to five cold exposures of −10 °C for 12 hrs and assessed 24 hours after).

Cold exposure and generation had a significant effect on energy reserves and metabolites measured in our biochemical assays. Cold exposure caused no significant differences in glycerol content (treatment: F_(3,34)_=1.039, p=0.39). This can be attributed to the large variance (sd=0.08) compared to the other treatment groups in F_0_ (sd<0.01; Fig. 6A). Glycogen content significantly differed among treatments (treatment: F_(3,34)_=7.60, p<0.001), with control and single 1 being significantly lower than single 2 and repeatedly cold exposed larvae (Fig. 6B). There was no significant interaction between treatment and protein mass (treatment × total protein: F_(3,34)_=2.03 p=0.13). Glycogen concentrations differed with protein mass (total protein: F_(1,34)_=43.81, p<0.001). Total carbohydrate significantly differed among treatments (treatment: F_(3,34)_=5.42, p<0.01). Post-hoc tests indicated larvae in treatments single 2 and repeated had significantly higher total carbohydrate content than larvae in the control and single 1 treatments (Fig. A1).

**Fig. 6.**
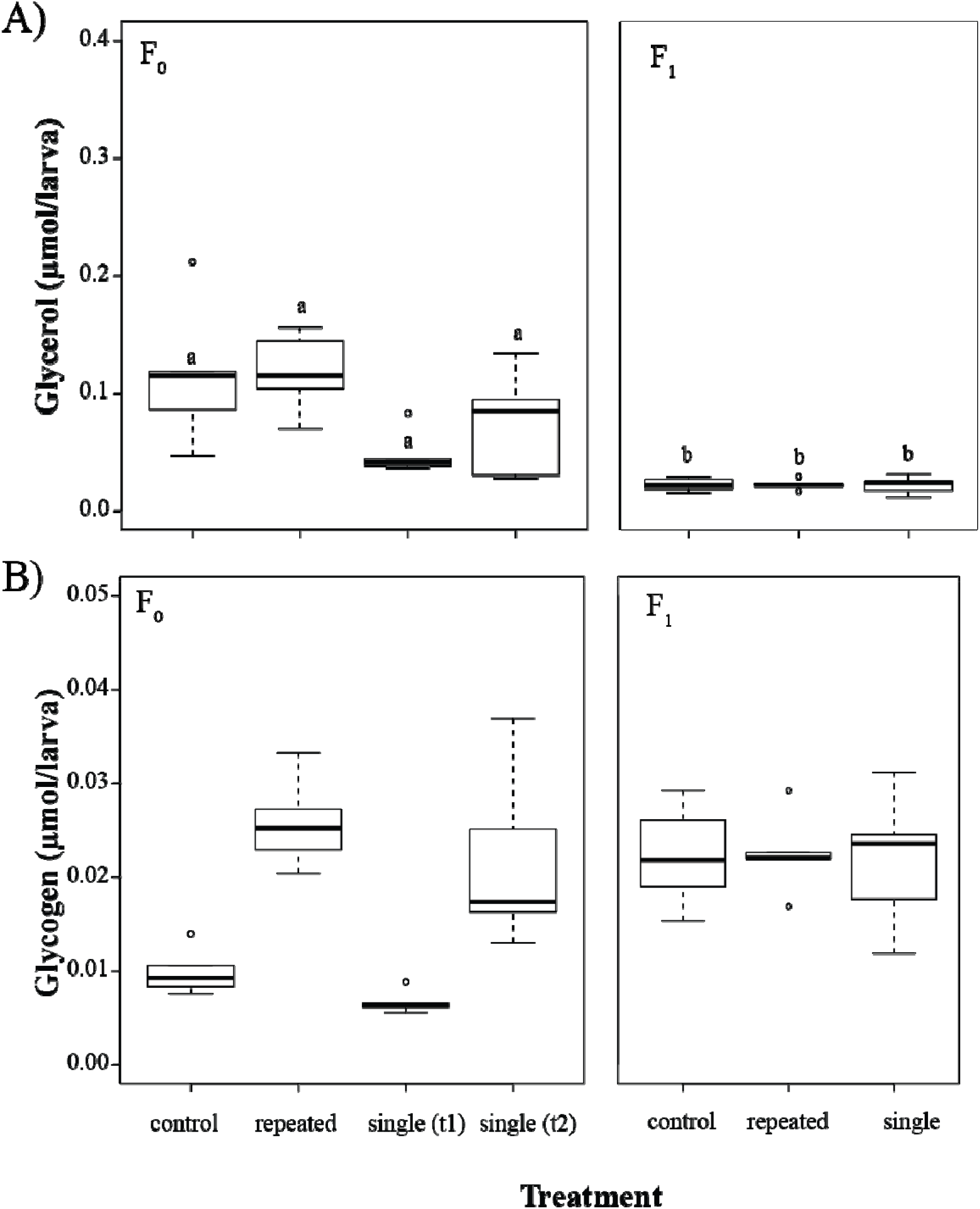
A) Glycerol per larva (μmol) and B) Glycogen per larva (μmol) across the different treatments and two generations in second-instar IPQL *Choristoneura fumiferana* larvae. Refer to Fig. 2 caption for explanation of boxplot display and treatment groups. Letters indicate statistically significant comparisons (p≤α). Glycogen is measured in glucose units. Points with the same letter are not statistically significantly different in an ANCOVA with protein mass as a covariate.

## Discussion

Geographic differences in cold tolerance can arise from genetic variation, phenotypic plasticity, or their interaction (reviewed in Chevin et al., 2010). Here, we explored the relationship between local adaptation and plasticity in overwintering capacity using *C. fumiferana*, an ecologically and economically important forest insect pest. We found regional differences in both seasonal and short-term plasticity of cold hardiness in *C. fumiferana*, suggesting local adaptation to winter climates. Regional differences in short-term plasticity were greater than differences in basal cold hardiness in early diapause larvae, emphasizing the importance of inducible responses in high latitude populations. We found no evidence for TGP despite a significant reduction in F0 fitness following repeated cold exposures. Our results highlight the importance of regional differences in cold tolerance, particularly the capacity for northern populations to respond to variable winter conditions. We suggest that future studies consider regional differences in spruce budworm populations and include seasonal and short-term plasticity when quantifying overwintering capacity.

Seasonal plasticity in insect cold tolerance is well described (Bale, 2002). Generally, our populations reduced their SCPs and shifted metabolite concentration as diapause progressed. Similar results were previously found in the IPQL laboratory strain, with cold tolerance and metabolites changing over the dormancy period (Marshall and Sinclair, 2015). However, our results indicated that these patterns of seasonal plasticity varied among populations. While our populations did not differ in their basal SCP in early diapause (Fig. 2A, 3A), late diapause larvae from high latitudes had significantly lower SCPs compared to lower latitude populations (Fig. 2A). These larvae also had significantly higher glycerol content compared to lower latitude populations and their increased glycerol content corresponded to reductions in SCP. Our results suggest that *C. fumiferana* has significant local adaptation to seasonal winter conditions.

High latitude populations experience significantly longer and colder winters than populations at lower latitudes (Marshall et al., 2020). Recent studies have shown that local adaptation in the degree of seasonal cold tolerance occurs across a range of native and invasive insects (Batz et al., 2020; Marshall et al., 2020; Williams et al., 2015). As expected, spruce budworm populations from high latitudes (e.g. Inuvik, Northwest Territories and High Level, Alberta) had greater overwintering capacity than eastern populations at lower latitudes. The basal SCP for late diapause Inuvik insects is lower than results published for *C. fumiferana* larvae from the IPQL lab colony under identical experimental and rearing conditions (Han and Bauce, 1995a; Marshall and Sinclair, 2015). In fact, we actually encountered methodological challenges measuring SCPs for these northern individuals. We were unable to freeze late-diapause Inuvik larvae using a refrigerated circulating bath, a method successful for other populations in our study. Therefore, we had to use a −80 °C freezer to reach temperatures low enough to initiate freezing in these larvae. An important difference in these two methods is that the cooling rate in the freezer was faster (9 °C/min) than the programmable bath (0.09 °C/min). There is evidence that cooling rates can influence absolute thermal limits (Terblanche et al., 2007), so the faster cooling rate of the freezer could have artificially depressed the SCP of the Inuvik insects. However, given that we switched to this method because the Inuvik population would not freeze under conditions used for other populations, and they consistently had lower SCP measurements than southern populations in other experiments (i.e. following repeated cold exposures, Fig. 6A), we are confident that the lower SCP in late diapause larvae is accurate.

Short-term plasticity allows organisms to respond rapidly to variable environmental conditions through inducible responses. We verified that fluctuating temperatures induced greater cold tolerance in *C. fumiferana* than constant temperatures alone (Marshall and Sinclair, 2015), and then demonstrated that high latitude populations expressed a significantly greater magnitude of plasticity than lower latitude populations (Fig 3). Short-term plasticity allows insects to respond to variable temperatures in a short temporal timeframe (reviewed in Marshall and Sinclair, 2012). The ability to rapidly respond to temperature shifts would be advantageous in northern climates where winter conditions are significantly more variable than southern latitudes (Marshall et al., 2020). Both seasonal and short-term plasticity combine to provide high latitude populations greater overwintering capacity than their southern counterparts.

Theoretical models often pit local adaptation against phenotypic plasticity (e.g. Sultan and Spencer, 2002), however these results indicate that phenotypic plasticity itself may be under local adaptation, particularly in insect overwintering. Spruce budworm populations in our study represented two distinct genetic backgrounds: Central and Eastern subclusters (Lumley et al., 2020). Our high latitude populations belong to the Central cluster, while New Brunswick and Quebec belong within the Eastern cluster. In fact, the IPQL population was founded from populations in eastern Canada, giving it the same genetic background as our other eastern populations. Even though the IPQL strain has been maintained in colony since the 1960s (Roe et al., 2017) and does not experience cold temperature stress, it was functionally equivalent to Eastern populations. Genomic differences between these central and eastern *C. fumiferana* populations are defined by a block of single nucleotide polymorphisms located on a single linkage group (LG4, Lumley et al., 2020). The authors noted that one of these variant loci was associated with glycerol-3-phosphate dehydrogenase (GPDH). GPDH is known to be involved in the glycerol synthesis pathway in other insects (Park and Kim, 2013), which is a key cryoprotectant in *C. fumiferana* (Han and Bauce, 1995b; Marshall and Sinclair, 2015) and insects in general (Lee, 2010). Larvae from the Central cluster produced elevated glycerol concentrations, so this genomic variation could be linked to the functional differences we observed. Further exploration of these genomic variants in the context of glycerol production will be critical to resolving the underlying mechanisms driving local adaptation in *C. fumiferana*.

Transgenerational plasticity (TGP) allows insects to survive in variable climates by inducing increased tolerance to temperature stress in future generations. First, however, we needed to demonstrate that repeated cold exposures triggered a cold stress response in the F0 generation. While we found that SCP did not change among the F0 treatment groups, we noted a decrease in LT_50_ in caterpillars that received repeated cold exposures. As such, we took this as a response to temperature stress and the insects altered their cold hardiness. Additionally, we found that treatment groups differed in mortality over time, suggesting that cold temperatures caused injury that manifested at later developmental stages. Although we did not observe a significant increase in glycerol content as a result of cold exposure, it is possible that unmeasured cold hardiness mechanisms (e.g. heat shock proteins or antifreeze proteins) could be responsible for this increased cold hardiness. Our results align with previous work (Han and Bauce, 1995a) and although tolerance to acute cold temperatures is very high as measured by SCP, the resultant chilling injury resulting from temperatures above the SCP is still significant. These results suggest that we induced a significant cold stress response in our F0 generation and established the selective environment for a transgenerational response.

Despite inducing a cold stress response in our F0 generation, we did not detect evidence of TGP in cold tolerance in our F1 generation. This lack of TGP after repeated cold exposures could be due to post-diapause feeding which masked the effect of stress-induced TGP (Tauber and Tauber, 1986). We did, however, find that F0 parents who experienced repeated cold exposures experienced a substantial decrease in fecundity (>50%), similar to results observed in the vinegar fly *Drosophila melanogaster* (Marshall and Sinclair, 2010) even when food was not limiting. Variable, stressful environments can impact many fitness traits, including growth, body size, energy stores, and development, ultimately leading to reduced fecundity (Buckley et al., 2021). High latitude or high elevation populations are exposed to highly variable temperature environments, which can lead to a combination of short-term mortality and long-term reproductive effects that impact overall population demographics (Buckley et al., 2021). Mechanistically, this lowered reproductive output could be due to an inability to replenish depleted energetic reserves or long-term damage. We did not detect differences in pupal weights, which suggests that individuals were likely able to replenish themselves during the post-stress feeding stage, and instead it is more likely that chronic damage was incurred that was not reflected in overall weight. Larval exposure to heat stress in *D. melanogaster* can cause developmental defects in eclosing adults which is ameliorated in strains with higher HSP70 copy number (and therefore expression), suggesting that unrepaired protein damage may persist through metamorphosis (Roberts and Feder, 1999). It is, therefore, not inconceivable that the same is true for cold stress in overwintering larvae (reviewed in King and MacRae, 2015), and would be worthwhile exploring in greater detail.

TGP can only be adaptive when environmental conditions are predictable between parent and offspring lifecycles. Larval *C. fumiferana* are short-distance dispersers; they are capable of dispersing between a few trees by “ballooning” on silken threads or walking to different parts of the tree crown (Johns and Eveleigh, 2013; Nealis, 2016). By comparison, adult moths disperse much larger distances, sometimes hundreds of kilometers (Sturtevant et al., 2013). Therefore, the probability of shared environmental conditions between generations is low, and TGP of cold hardiness in second-instars is not favored by selection. It is also possible that TGP occurs in other winter-related traits rather than in absolute cold hardiness or supplied energetic reserves. For example, Harvey (1961) found that some fourth-instar *C. fumiferana* undergo a second diapause. The expression of a second diapause may be a plastic response to environmental variability. If so, TGP could be observed in diapause expression among larval instars since it is likely that environmental predictability between instars is higher than between generations. The lack of TGP in our measures of cold hardiness could also be due to using a lab-selected strain. The IPQL strain has been in culture since 1961 and has not been augmented with wild material for the past 20 years (Roe et al., 2017). Therefore, the relaxed selection of cold hardiness under lab conditions could have resulted in a very different response compared to wild populations. Wild populations may differ in this respect, and would be an important area for further study.

## Conclusions

The eastern spruce budworm has a broad geographic range in North America, so populations are exposed to very different temperature regimes across this range. It was hypothesized that populations would exhibit local adaptation in seasonal as well as short-term plasticity. Two populations tested, Alberta and Inuvik, showed a similar degree of short-term plasticity, which we attribute to the geographic connectivity between the populations. Generally, there were no differences in basal cold hardiness among the populations we tested, however seasonal and repeated cold exposures revealed evidence for local adaptation of plasticity in the populations. These results have direct implications for the predicted population growth, range shifts, and current species distribution modelling for the species. The IPQL lab strain, which forms the basis of our current understanding of cold hardiness in the species, shows comparatively low plasticity. Therefore, shifting thermal regimes in local environments due to climate change, leading to increased temperature extremes that select for plasticity, will result in highly variable regional responses among spruce budworm populations.

Winter and higher latitudes have experienced the most change in temperature under new climate conditions (Marshall et al., 2020; Zhang et al., 2019), exposing populations to novel thermal environments across their ranges. Understanding the regional responses of populations across a species’ range is critical for predicting future changes in population growth and species distribution throughout the boreal forest. Absence of evidence for cold hardiness TGP in the IPQL strain suggests that TGP does not need to be incorporated into population growth or species range models in *C. fumiferana*. However, it does provide further evidence that repeated cold exposure may drive population growth effects and that fitness trade-offs in overwintering insects exist. Further work should focus on untangling the potential mechanisms of these trade-offs, and future modelling should include fitness effects of repeated cold exposures.

Local adaptation and phenotypic plasticity can and should be included in species distribution models for *C. fumiferana*. Bush et al. (2016) highlights the use of hybrid species distribution models, which uses trait mean, variability, heritability and the plasticity of a trait to determine range shifts, therefore incorporating both plasticity and evolution. Diamond (2018) tested this *AdaptR* model and found that range loss for 17 species of *Drosophila* decreased by 33% in 2105 when incorporating these additional traits. For *C. fumiferana*, accurately modelling species distributions with these spatio-temporal adaptive models could prove useful for species management in future climates.

## Supporting information

Supplemental results

## Acknowledgments

The authors with to express gratitude to everyone who helped with the creation of this manuscript. Thanks to Ashlyn Wardlaw, Kerry Perrault, and the Insect Production and Quarantine Laboratories (Natural Resources Canada) for rearing and maintaining the spruce budworm colonies central to this project.

## Author Contributions

Conceptualization, S.B., A.D.R. and K.E.M.; laboratory experiments, S. B.; analyses, S.B. and K.E.M.; manuscript preparation, S.B., K.E.M.

## Data availability statement

Data and code is available on the Open Science Framework. DOI: 10.17605/OSF.IO/ZNEFB

