## Supplemental results for "Plasticity of cold hardiness in the eastern spruce budworm, *Choristoneura fumiferana*"

Supplementary data.

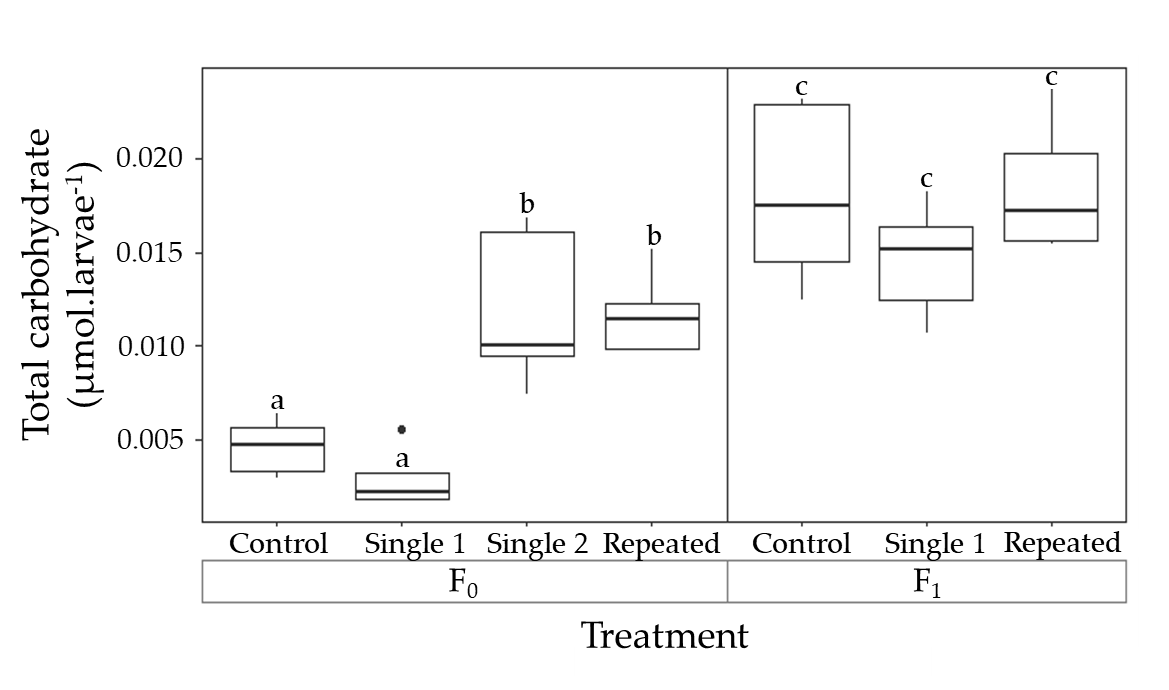

Fig. A1. Total carbohydrate per larva (μmol) across the different treatments and two generations in second-instar IPQL *Choristoneura fumiferana* larvae. Refer to Fig. 2 caption for explanation of boxplot display. Treatment groups are Single 1 (larvae exposed to a single cold exposure of -10 °C for 12 hrs and assessed 24 hours after), Single 2 (larvae exposed to a single cold exposure of -10 °C for 12 hrs and assessed at the same time as the repeated group) and Repeated (larvae exposed to five cold exposures of -10 °C for12 hrs and assessed 24 hours after). Different letters indicate statistically significant comparisons (p≤α).

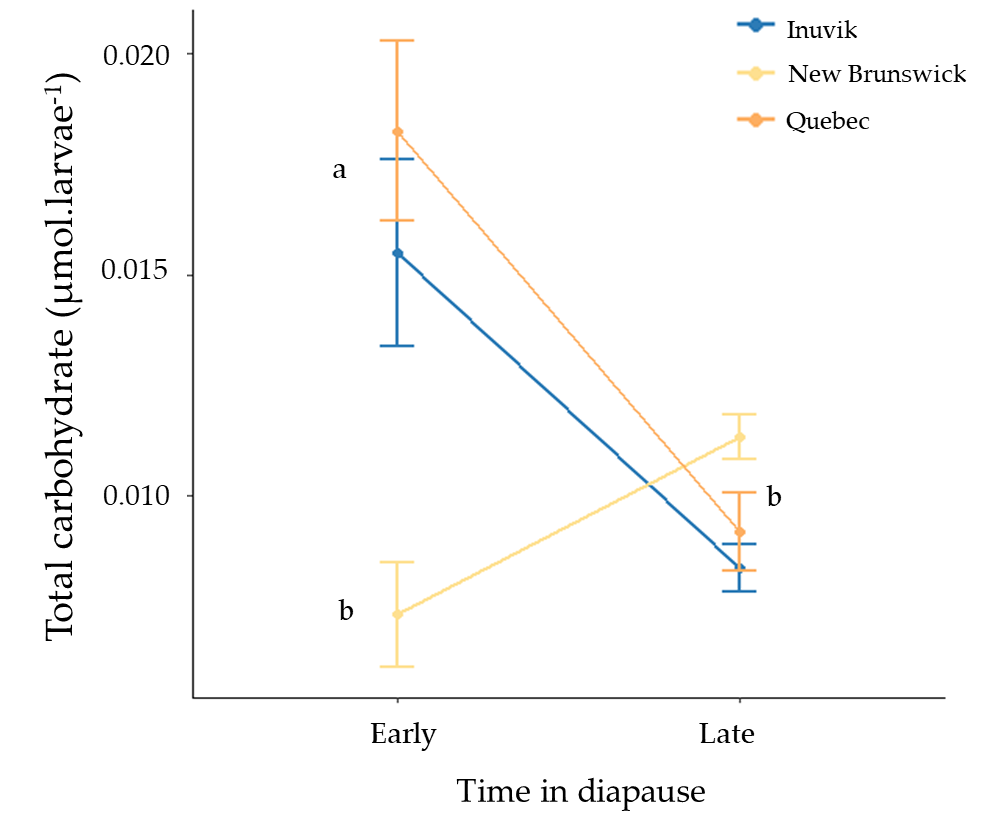

Fig. A2. Basal total carbohydrate per larva (μmol) of second-instar *Choristoneura fumiferana* across different populations tested at either early (6 weeks) or late (12 weeks) into diapause. Different letters indicate statistically significant comparisons (p≤α).

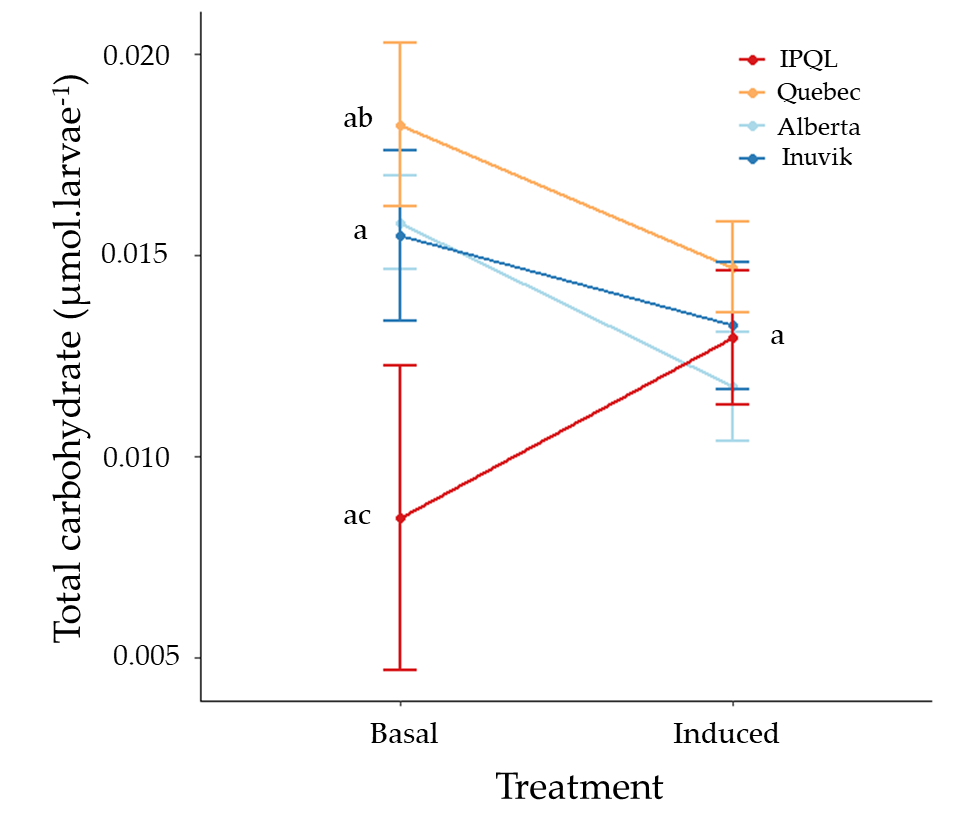

Fig. A3. Total carbohydrate (μmol) of second-instar *Choristoneura fumiferana* before (“basal”) and after (“induced”) five exposures to -15 °C. Different letters indicate statistically significant comparisons (p≤α).

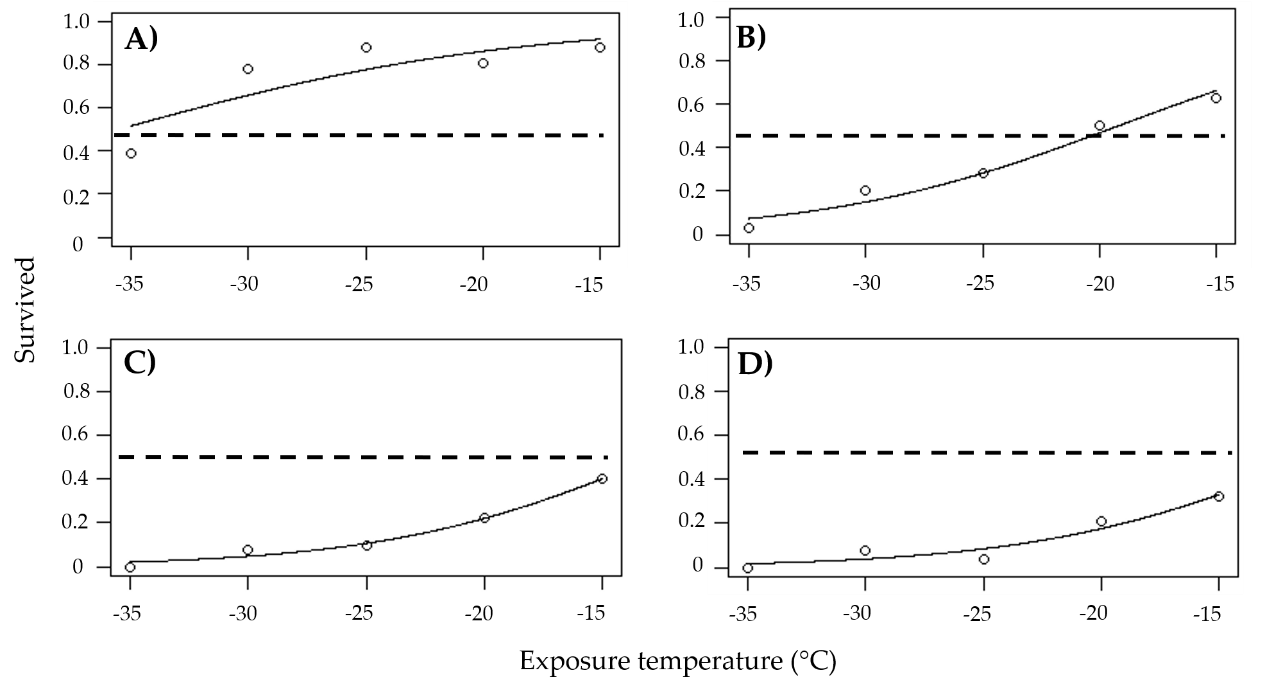

Fig. A4. Proportion of survival of Control F_0_ second-instar IPQL (*Choristoneura fumiferana*) larvae assessed at different developmental times after 4-hour exposures to lower lethal temperature treatments (n=100 per treatment). Times assessed are A) 1 week, B) the end of diapause, C) thinning, and D) pupation.

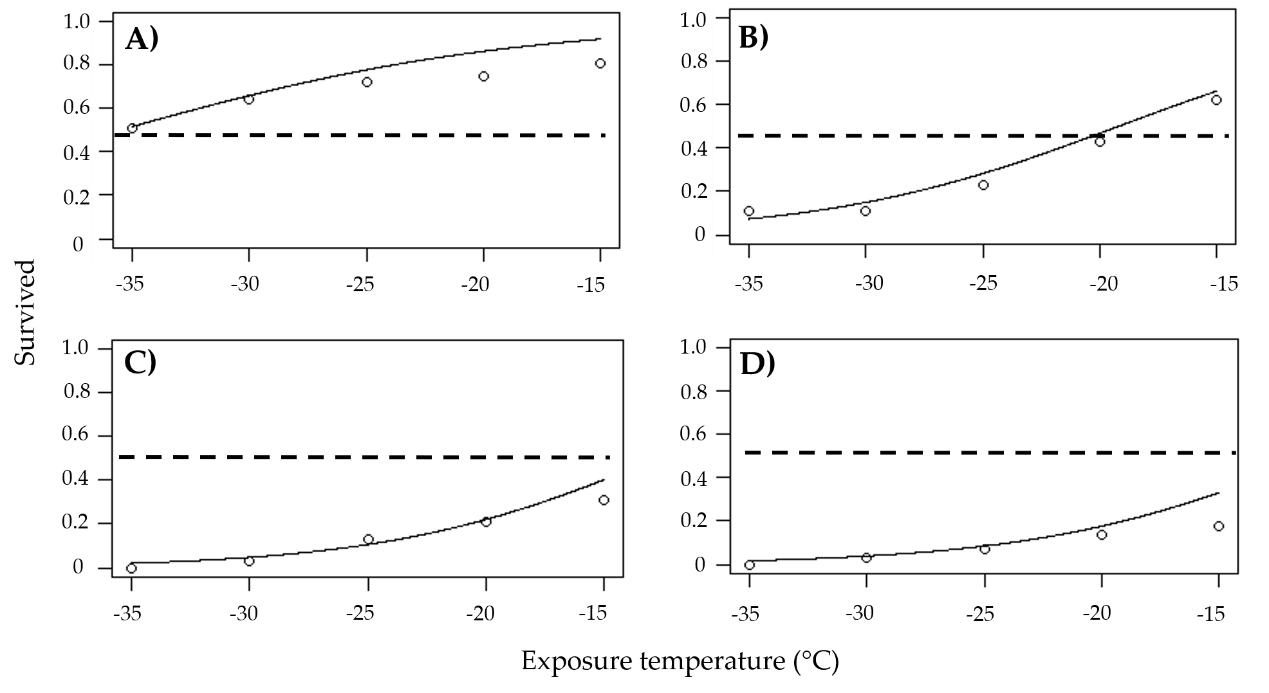

Fig. A5. Proportion of survival of Single 1 F_0_ second-instar IPQL (*Choristoneura fumiferana*) larvae assessed at different developmental times after 4-hour exposures to lower lethal temperature treatments (n=100 per treatment). Times assessed are A) 1 week, B) the end of diapause, C) thinning, and D) pupation.

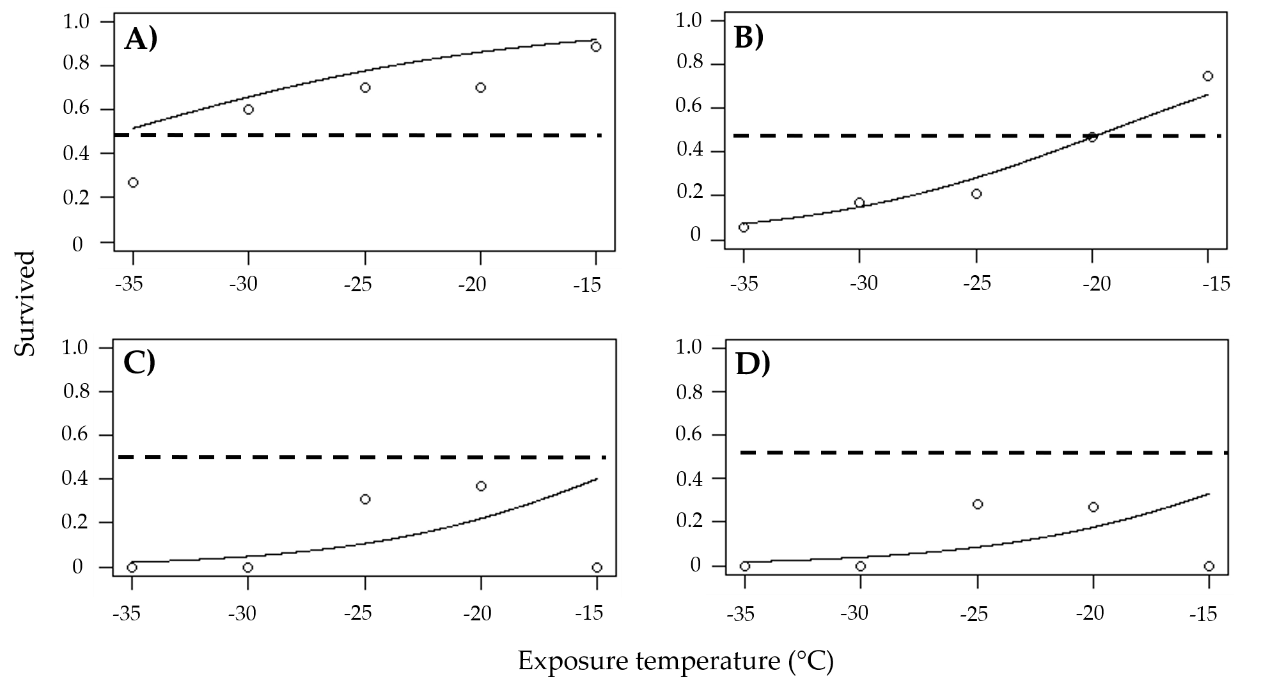

Fig. A6. Proportion of survival of Single 2 F_0_ second-instar IPQL (*Choristoneura fumiferana*) larvae assessed at different developmental times after 4-hour exposures to lower lethal temperature treatments (n=100 per treatment). Times assessed are A) 1 week, B) the end of diapause, C) thinning, and D) pupation.

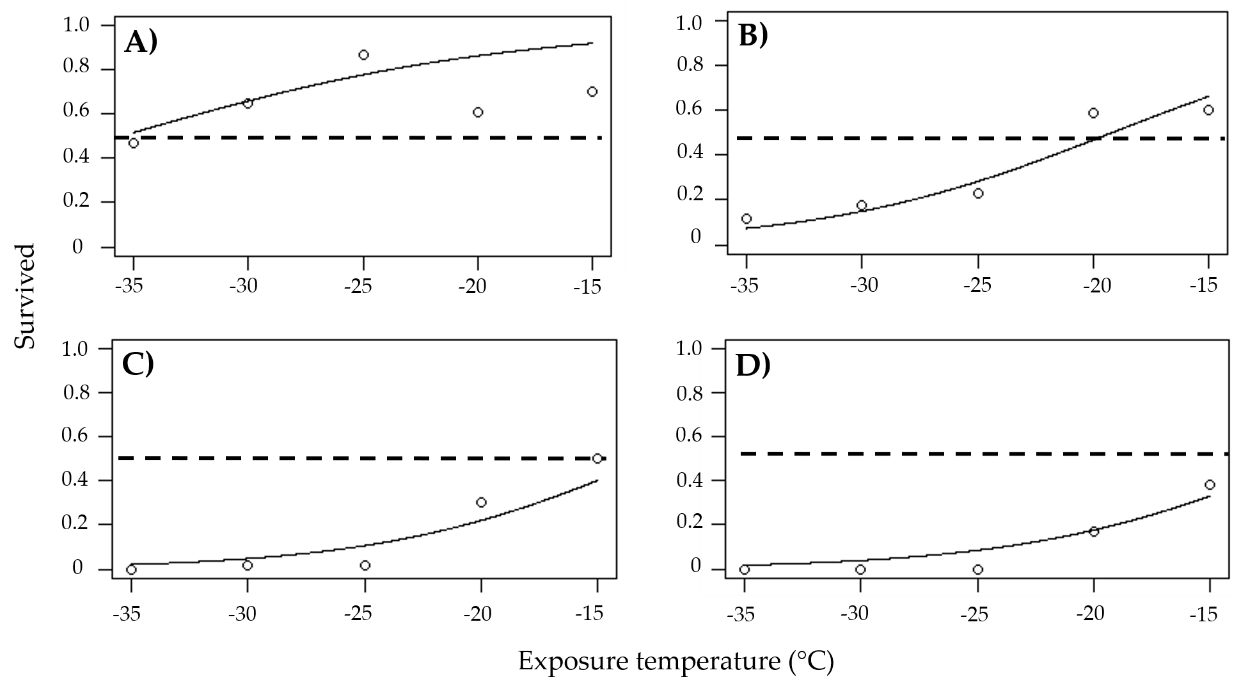

Fig. A7. Proportion of survival of Repeated F_0_ second-instar IPQL (*Choristoneura fumiferana*) larvae assessed at different developmental times after 4-hour exposures to lower lethal temperature treatments (n=100 per treatment). Times assessed are A) 1 week, B) the end of diapause, C) thinning, and D) pupation.
